# Positive association between alleles at selectively neutral loci

**DOI:** 10.1101/2023.06.14.544962

**Authors:** Nadezhda A. Potapova, Alexey S. Kondrashov

## Abstract

Linkage disequilibrium (LD) is a non-independent distribution of alleles at different loci and might be defined using directional and directionless approaches. While analytical theory is available for directionless LD, analytical theory for directional LD is less developed. In this study we investigated directional LD generated by random drift between two selectively neutral loci. When the sign of LD is determined by the ancestral-derived status of these alleles, its expected value is zero. In contrast, computer simulations show that random drift leads to a tendency of rare, as well as of common, alleles to occur together. This observation is supported by a straightforward analytical argument. If only those generations where both loci are polymorphic are taken into account, the magnitude of the resulting directional LD depends only on N_e_. These observations might give an opportunity to directly estimate N_e_ from the data on genetic variation within a single generation.

## INTRODUCTION

Non-independent distribution of alleles at different loci, referred to as linkage disequilibrium (LD) between them, is one of the key phenomena studied by population genetics. In the simplest case of two haploid loci A (with alleles A and a) and B (with alleles B and b), LD is characterized by D = [ab] - [a][b] (Lewontin and Kojima 1960), where square brackets denote frequencies of the genotypes carrying both alleles a and b, only a and only b, respectively. D, as well as its normalized derivatives such as D’ or r (see Charlesworth and Charlesworth 2009), is a directional measure of LD, in the sense that its sign depends on the direction of association between the two loci: it is positive if alleles a and b, as well as A and B, tend to occur together, and negative if they avoid each other. As long as these two situations are equally common, which must be the case when alleles are denoted by capital and lowercase letters in an arbitrary way, the expected value of a directional measure of LD is zero. Then, it makes sense to characterize LD by a directionless measure, such as r^2^. And, of course, non-zero directionless LD is consistent with zero directional LD.

Still, there is a number of situations when alleles A and a, and B and b, have different properties, so that alleles a and b tend to either occur together or to avoid each other. The most obvious case of this kind is epistatic selection: alleles whose combination confers a lower fitness than the one expected, if they acted independently, avoid each other (Langley and Crow 1974, Boyrie et al. 2021). If so, it makes sense to characterize LD by a directional measure.

However, the leading force responsible for LD in amphimictic populations is not selection but random drift (Hill and Robertson 1968, Ohta and Kimura 1969). Indeed, alleles at loci that are physically distant from each other are distributed mostly independently, so that LD of any kind between them is close to zero. In contrast, when recombination is slow, patterns of directionless LD between selectively neutral sites are not radically different from that between sites that affect fitness (*e. g*., Kim et al. 2007), because of random drift.

Can random drift generate directional LD? The answer to this question may depend on how we polarize genetic variation at a neutral diallelic locus. There are at least two natural ways of doing so. First, we can regard as positive coupling associations between rare (and, thus, between common) alleles (eq. 1 in Langley and Crow 1974; Pool et al 2012; Ragsdale 2022; Good 2022). Second, we can regard as positive coupling associations between young, *i. e*. derived (and, thus, between old, *i. e*. ancestral) alleles (Takahasi and Innan 2008).

In contrast to the case of directionless LD, where analytical theory is available (*e*.*g*. Hill and Robertson 1968; Ohta and Kimura 1969), there is no such developed general analytical theory for directional LD. However, there are many observations on directional LD obtained mostly using population datasets. For instance, Pool et al. (2012) observed both positive and negative associations between rare alleles in a natural population of *Drosophila melanogaster* (Fig. 11 in Pool et al. 2012), but did not address the issue of the average sign of these associations. Sandler et al. (2021, Fig. 3) investigated two population genomic datasets (*Capsella grandiflora* and *Drosophila melanogaster*) and reported that rare synonymous and intron alleles tend to occur together. Ragsdale (2022) showed on computer simulations and human population dataset that there is a positive LD between synonymous mutations, but did not discuss the role of a random drift separately. Examination of data from 1000 Genomes Project as well as computer models showed that there is a lower LD value for derived nonsynonymous alleles than for synonymous alleles (Garcia and Lohmueller 2021). Using a computer simulation, Takahasi and Innan (2008) showed that on average signed LD between neutral loci polarized by their age is zero.

In this study we use computer simulations to demonstrate that random drift generates positive LD between neutral loci polarized by allele frequency, providing a heuristic explanation for this effect. We also confirm and explain the result of Takahasi and Innan (2008) by a straightforward analytical argument.

## MATERIAL AND METHODS

We simulated a haploid Wright-Fisher population polymorphic at two linked diallelic loci A and B with bidirectional mutation and studied directional and directionless LD between them under different values of the population size N, mutation rate m (rates of mutation in both directions were assumed to be equal), rate of recombination r, and coefficient of selection s against alleles a and b. Each entry was the average of 20 runs, each recorded in the course of 50 million generations, after a burn-in of 1 million generations. The model was implemented as a C code, which is available upon request.

Directional LD was characterized by

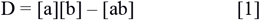

An allele was regarded as “lowercase” either if it was rare (having frequency below 0.5) or if it was derived. We separately report the average value of D for all generations and only for “polymorphic” generations, *i. e*. those generations when both loci A and B were polymorphic in the simulated population, making a non-trivial LD possible. Directional LD was characterized by

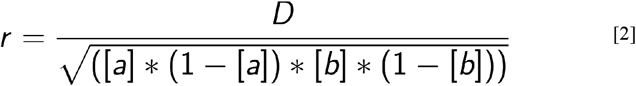

(see Charlesworth and Charlesworth 2009), and directionless LD was characterized by r^2^.

For all simulations computational resources of the “Makarich” HPC cluster were provided by the Faculty of Bioengineering and Bioinformatics, Lomonosov Moscow State University, Moscow, Russia.

## RESULTS

### Positive associations between rare neutral alleles

First, we studied a population with no recombination or selection, i. e. with r = 0 and s = 0. Such a population is fully described by its actual size N, which is also its effective size N_e_, and the mutation rate m. Measures of LD recorded for all generations and only for polymorphic generations were denoted with subscripts “all” and “poly”, respectively. “r(age)” and “r(freq)” are measures of directional LD polarized by the allele ages and frequencies, respectively.

In Table 1 we present data on LD obtained for m = 10^−5^. If only polymorphic generations are taken into account, directionless LD, characterized by r^2^_poly_, decreases when N increases, because it is generated by random drift. In contrast, when all generations are taken into account, directionless LD r^2^_all_ increases when N increases, because when Nm is small the probability that a generation is polymorphic is proportional to (Nm)^2^, so that when N is low, for a vast majority of generations LD is trivially equal to zero.

**Table 1.** The dependence of directionless LD and directional LD polarized by allele ages and frequencies on the population size. Standard errors are shown in parentheses.

| Population size | $r^2_{\text{all}}$ | $r^2_{\text{poly}}$ | $r(\text{age})_{\text{all}}$ | $r(\text{age})_{\text{poly}}$ | $r(\text{freq})_{\text{all}}$ | $r(\text{freq})_{\text{poly}}$ |
| --- | --- | --- | --- | --- | --- | --- |
| 100 | 0.000018<br>(0.000002) | 0.190454<br>(0.012281) | 0.000002<br>(0.000002) | 0.014971<br>(0.018731) | 0.000012<br>(0.000002) | 0.123050<br>(0.014209) |
| 1 000 | 0.001697<br>(0.000055) | 0.095618<br>(0.002649) | 0.000060<br>(0.000048) | 0.003392<br>(0.002738) | 0.001116<br>(0.000054) | 0.062865<br>(0.002769) |
| 10 000 | 0.056185<br>(0.000567) | 0.080664<br>(0.000767) | 0.000078<br>(0.000862) | 0.000124<br>(0.001237) | 0.033619<br>(0.000609) | 0.048266<br>(0.000854) |
| 100 000 | 0.000005<br>(0.000235) | 0.000005<br>(0.000235) | -0.002375<br>(0.005916) | -0.002375<br>(0.005916) | 0.039046<br>(0.003009) | 0.0390456<br>(0.003009) |

In agreement with Takahata and Innan (2008) directional LD polarized by allele age is not significantly different from 0. In contrast, we observed that random drift generates systematic attraction between rare (and, thus, also between common) alleles, leading to positive values of r(freq). As it is the case for directionless LD, this association is the strongest with low N if only polymorphic generations are taken into account (r(freq)_poly_), and for high N if all generations are taken into account (r(freq)_all_). When N increases, the proportion of polymorphic generations also increases, and with N = 100 000, all generations are polymorphic, leading to the same values of r(freq)_poly_ and r(freq)_all_.

### LD in polymorphic generations is independent of the mutation rate

Table 2 shows how r(freq)_poly_ and r(freq)_all_ depend on the mutation rate with N = 1000. As we can see, r(freq)_poly_ is independent of the mutation rate. This is not surprising, because, under a low Nm, the population is polymorphic at both loci only in the course of rare, isolated episodes, and during each such episode no new mutations occur, so that the dynamics of the genetic composition in the course of an episode does not depend on the mutation rate. In contrast, r(freq)_all_ is proportional to m^2^, because in the absence of selection the fraction of generations when both loci are polymorphic is proportional to m^2^.

**Table 2.** The dependence of directional LD with alleles polarized by their frequencies on the mutation rate. Standard errors are shown in parentheses.

| Mutation rate | $r(\text{freq})_{\text{all}}$ | $r(\text{freq})_{\text{poly}}$ |
| --- | --- | --- |
| $10^{-4}$ | 0.034071<br>(0.000227) | 0.062430<br>(0.000397) |
| $10^{-5}$ | 0.001117<br>(0.000054) | 0.062865<br>(0.002769) |
| $10^{-6}$ | 0.000013<br>(0.000005) | 0.049510<br>(0.015933) |

### LD rapidly disappears when recombination increases

Table 3 presents data on the dependence of directionless and directional LD polarized by allele frequency on recombination rate c, under N = 10 000, m = 10^−5^, and s = 0. As expected, even weak recombination is sufficient to effectively destroy LD generated by random drift (Hill and Robertson, 1968).

**Table 3.** The dependence of LD on the rate of recombination. Standard errors are shown in parentheses.

| Recombination rate | $r^2(\text{freq})_{\text{all}}$ | $r^2(\text{freq})_{\text{poly}}$ | $r(\text{freq})_{\text{all}}$ | $r(\text{freq})_{\text{poly}}$ |
| --- | --- | --- | --- | --- |
| $10^{-4}$ | 0.056185<br>(0.000567) | 0.080664<br>(0.000768) | 0.033619<br>(0.000609) | 0.048266<br>(0.000854) |
| $10^{-5}$ | 0.015868<br>(0.000094) | 0.022781<br>(0.000119) | -0.001593<br>(0.000155) | -0.002286<br>(0.000222) |

### LD decreases under constant negative selection

Table 4 presents data on the dependence of directionless and directional LD polarized by allele frequency on the strength of negative selection, *i. e*. the coefficient of selection s, under N = 10 000, m = 10^−5^, and c = 0. We can see that when s increases, both these characteristics rapidly decline, with directional LD declining faster than directionless LD.

**Table 4.**
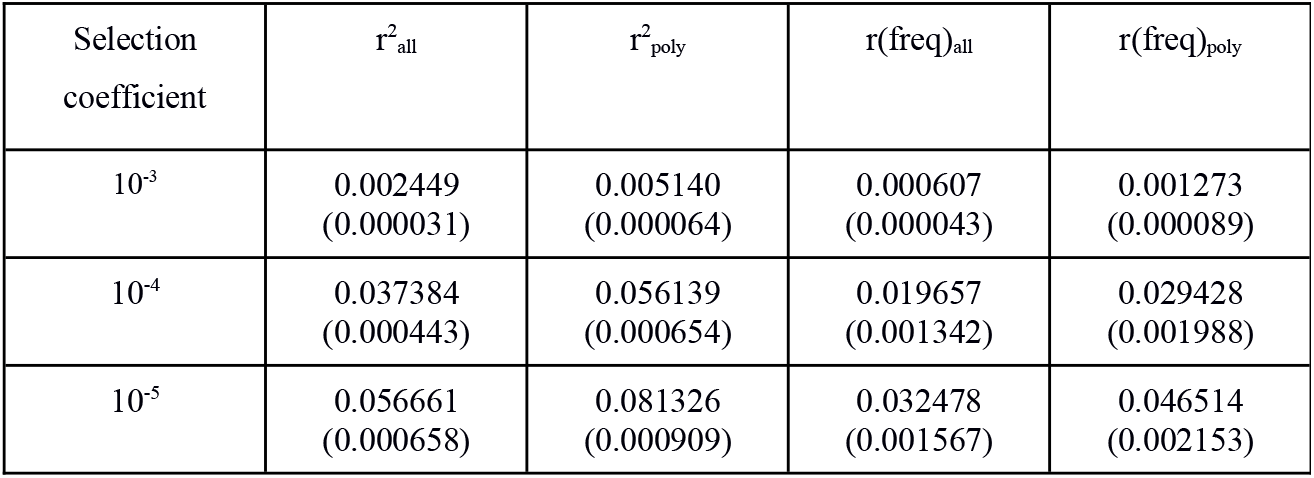
The dependence of characteristics of LD on the selection coefficient. Standard errors are shown in parentheses.

### Expected LD between neutral derived alleles is zero

Let us first demonstrate that the expected LD is zero for a newly arising variant pairs. A new pair arises when one mutation occurs in first loci (say, in locus A, that segregates in the population with alleles A and) and another mutation – in a second locus (say, locus B with alleles B and b). If this mutation, b, occurs on the background of the derived allele in locus A, i.e. loci a, then the LD will be is as mentioned in formula [1] in this study.

If the new mutation at locus B, i.e. b, occurs on the background of the ancestral allele at locus A, i.e. allele A, LD is -[ab]. The probability of the new allele at locus B to arise on the background of the derived allele at locus A is (1-[a]).

Finally, the expected value of D will be

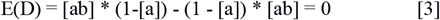

Therefore, in the absence of recombination, selection and random genetic drift, considering only mutations, we expect no LD between two alleles.

Now, let’s consider a change in LD due to drift. In case when N_e_m << 1 we have 3 frequencies [ab], [Ab], [aB] for 3 haplotypes (numbers of individuals of each haplotype are N1, N2 and N3, respectively). In the absence of recombination, LD is a product of the frequencies of 2 haplotypes (either positive or negative) and we need to look at the population sample in the next generation which is a sample from the trinomial distribution:

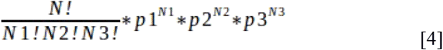

Expected LD in the next generation is given by:

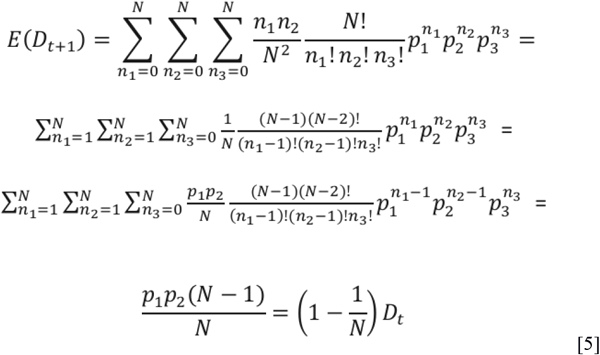

For a diploid population, E(Dt+1)=(1-1/2N)*D_t_ [6]. Now, we see that the expected D is zero at the moment of origin and changes by a constant multiplicative factor at each generation. This means that the overall expected LD is zero.

## DISCUSSION

The only one obvious reason of non-zero LD between loci is epistatic selection. Indeed, if effect of combination of loci is more radical than it could be expected and decreases fitness dramatically, there is a place for epistatic selection which causes negative LD (Langley and Crow 1974). Therefore no LD might be expected between two selectively neutral loci.

Indeed, Takahasi and Innan (2008) using computer simulations showed zero directional LD between neutral alleles polarized by age. In our study we confirmed their result, but furthermore observed positive associations between alleles polarized by frequency.

The only explanation of positive LD between neutral alleles is random drift, that drives them to stay together. Evidence of directionless LD (r^2^) due to random drift is classical (Hill and Robertson 1968, Ohta and Kimura 1969), but existence of directional LD between neutral loci is a difficult issue and there is no general analytical theory for it. When we introduced non-zero selection coefficient in model – even weak selection almost completely destroyed the effect of attraction (Table 4). In such case rare alleles turned out equivalent to derive and become polarized by age rather than frequency. Indeed, when Ne*s > 1, the deleterious allele can never reach a high frequency, so that a derived allele always remains rare – and two polarizations, by allele age and by its frequency, become equivalent. Increased recombination and mutation rates also disrupt LD between pairs of loci (Table 2, 3).

Using straightforward theoretical arguments we explained why the overall expected LD between neutral derived loci is zero. But also it can be shown that the expected LD between minor alleles is positive. To develop intuition into this question, we can analyze trajectories of positively and negatively linked pairs of variants (in terms of derived alleles). If two variants are in negative LD for their derived alleles, for minor alleles LD remains negative for the part of trajectories where both derived alleles are below 50%. For the part of the trajectory where one of the variants is above 50% and the other is below 50%, LD between minor alleles becomes positive. Obviously, the situation where both derived alleles for negatively linked variants are above 50% is not possible.

For positively linked variants (in terms of derived alleles), LD remains positive for the part of the trajectory where frequencies of both variants are below 50%. LD polarized by frequency becomes negative where frequency of one variant is above 50% and frequency of the other variant is below 50%. Next, LD for minor alleles becomes positive again when both frequencies are above 50%. This effect results in the overall positive expectation for LD when polarized by allele frequency.

In the situation when both loci are polymorphic, LD value depends only on Ne. If Ne*m << 1, the strength of LD does not depend on mutation rate m, if only those generations when both loci are polymorphic are taken into account. This result is easy to explain: although the frequency of “polymorphic episodes” depends on the mutation rate, the dynamics of each such episode do not – and with Ne*m such episodes usually do not overlap with each other. As long as Ne*m is well below 1, so that “polymorphic episodes” are isolated, the magnitude of association between rare alleles depends only on Ne, creating an opportunity to estimate it independently of m. The possibility to directly calculate Ne by LD was previously shown by Hill (1981) who suggested to measure Ne using directionless r^2^, but we assume there is an opportunity to measure it from directional LD.

Overall, we observed positive associations between selectively neutral rare alleles that can be explained by random drift only. Such phenomena was clearly seen in computer simulations and supported by theoretical arguments.

## Acknowledgements

We thank Shamil R. Sunyaev for discussion and help in derivation of formulas. We thank Georgii A. Bazykin and Ksenia A. Khudyakova for helpful comments.

